# Adaptation of Plasmid-ID Technology for Evaluation of N^2^-Fixing Effectiveness and Competitiveness for Root Nodulation in the *Sinorhizobium-Medicago* System

**DOI:** 10.1101/2025.03.04.641427

**Authors:** Jake Schumacher, Nicholas Dusek, Marcela Mendoza-Suárez, Barney A. Geddes

**Affiliations:** Department of Microbiological Sciences, North Dakota State University, Fargo, ND, USA; Center for Computationally Assisted Science and Technology, North Dakota State University, Fargo, ND, USA; Department of Department of Molecular Biology and Genetics, Aarhus University, Denmark

**Author notes:** Corresponding Author: Barney Geddes.

**Keywords:** Synthetic biology, Symbiosis, Competition, Effectiveness, Sinorhizobium, Medicago, Alfalfa, Rhizobia, Biosensor, Barcode

## Abstract

Maximizing the nitrogen fixation occurring in rhizobia-legume associations represents an opportunity to sustainably reduce nitrogen fertilizer inputs in agriculture. High-throughput measurement of symbiotic traits has the potential to accelerate the identification of elite rhizobium/legume associations and enable novel research approaches. Plasmid-ID technology, recently deployed in *Rhizobium leguminosarum*, facilitates the concurrent assessment of rhizobium nitrogen-fixing effectiveness and competitiveness for root nodulation. This study adapts Plasmid-ID technology to function in *Sinorhizobium* species that are central models for studying rhizobium-legume associations and form economically important symbioses with alfalfa. New Sino-Plasmid-IDs were developed and tested for stability and their ability to measure competitiveness for root nodulation and nitrogen-fixing effectiveness. Rhizobial competitiveness is measured by identifying strain-specific nucleotide barcodes using Next-Generation Sequencing while effectiveness is measured by GFP fluorescence driven by the synthetic *nifH* promoter. Sino-Plasmid-IDs allow researchers to efficiently study competitiveness and effectiveness in a multitude of *Sinorhizobium* strains simultaneously.

## Introduction

Rhizobia and legume plants have a symbiotic relationship. Legumes host rhizobia in specialized organs called nodules developed on their roots and provide them with nutrients. In return, the rhizobia fix N_2_ from the atmosphere into reduced and bioavailable forms of nitrogen that are supplied to the legume (Sprent, 2007). This symbiotic relationship has been studied extensively due to the rhizobia’s utility as a substitute for chemical fertilizers in agriculture, either directly through the use of legume crops, or indirectly through crop rotation (Parnell *et al*., 2016). The inefficient overuse of nitrogen fertilizers in agriculture releases reactive nitrogen into the environment polluting the atmosphere, rivers, and coastal waters (Rockström *et al*., 2009) costing an estimated $210 billion a year in the United States alone (Sobota, 2015). Because of the detrimental impact of chemical fertilizers on the environment, there is an urgent need to maximize symbiotic nitrogen-fixation in agriculture. Long-term efforts to establish a nitrogen-fixing symbiosis in non-legume crops analogous to the legume-rhizobia-like symbiosis are currently underway (Mus *et al*., 2016). More immediately, maximizing the nitrogen fixation occurring in existing rhizobia-legume associations represents an opportunity to sustainably reduce nitrogen fertilizer inputs.

There is a large variance in both how effectively a rhizobium can fix nitrogen to benefit its host as well as their ability to compete to form a root nodule against native rhizobia found in the natural microbiome of a soil (Yates *et al*., 2011; Heath and Stinchcombe, 2014; Batstone *et al*., 2022). The “rhizobium competition problem” describes how rhizobium inoculants that are excellent at fixing nitrogen can be irrelevant to the fitness of a legume because they cannot form root nodules when competing against other strains of highly competitive rhizobia from the soil with inferior nitrogen-fixing ability (Triplett and Sadowsky, 1992; Yates *et al*., 2011; Mendoza-Suárez *et al*., 2021; Burghardt and diCenzo, 2023). Therefore, ideal rhizobial inoculants must be, first, highly competitive and, second, outstanding at nitrogen fixation. The model rhizobia *Sinorhizobium meliloti* is an alfalfa (*Medicago sativa*) and barrelclover (*Medicago truncatula*) symbiont. Due to its numerous genetic resources, including a deletion library (diCenzo *et al*., 2016), minimized genome (Geddes *et al*., 2021), and its development as a platform for population genomics (Epstein *et al*., 2018, 2023; Riley *et al*., 2023), *S. meliloti* is uniquely qualified to investigate genetic factors that underly key traits in the legume-rhizobia symbiosis. Furthermore, alfalfa is an important economic crop to the United States with the potential to provide all the nitrogen requirements for a subsequent crop (Yost *et al*., 2012), and *M. truncatula* is a model legume facilitating complementary genetic studies in the host (Roy *et al*., 2020). Another *Sinorhizobium* species, *Sinorhizobium medicae* is also an emerging model to contrast variance in symbiotic traits across species that share a similar host range (Liu *et al*., 2022; Yu *et al*., 2024). The genetically amenable model *S. medicae* WSM419 has been developed for its superior symbiotic performance with *M. truncatula* relative to *S. meliloti* model strains (Reeve *et al*., 2010).

Measuring rhizobium competitiveness typically involves low-throughput methods capable of comparing only a few strains at a time. Pairwise competition experiments have been routinely performed using the antibiotic resistance markers from insertion mutants, or *gusA* (stains magenta) and *celB* (stains blue) marker genes to compare the frequency of two strains in root nodules (Sánchez-Cañizares and Palacios, 2013; Geddes, González and Oresnik, 2014). More recently, next generation sequencing (NGS) has unlocked a variety of methods for more high-throughput assessments of rhizobium competitiveness based on shotgun or amplicon sequencing of nodule populations (Burghardt *et al*., 2018; Rahman *et al*., 2023). “Plasmid-IDs” are a recently developed technique that combines DNA barcoding of rhizobium strains for assessment of competitiveness by cost-effective amplicon sequencing with a *PnifH*:sfGFP biosensor that facilitates predictions of nitrogen-fixing effectiveness based on nodule fluorescence (Mendoza-Suárez *et al*., 2020, 2024). While this advancement is useful in rapidly characterizing rhizobial strains, the Plasmid-ID library constructed previously is optimized for *Rhizobium leguminosarum* and does not function in some important rhizobia such as *S. meliloti* (Mendoza-Suárez *et al*., 2020). In this study, we develop and validate a unique Plasmid-ID system for use in *Sinorhizobia*, allowing for the study of competitiveness and effectiveness in a multitude of *Sinorhizobium* strains simultaneously.

## Materials and Methods

### Bacterial growth conditions and media

Information on the *Escherichia coli*, *Sinorhizobium. sp.,* and *R. leguminosarum* strains used in this study can be found in Table 1. *E. coli* strains were grown on solid and liquid Luria-Bertani medium (LB) at 37°C. *Sinorhizobium* and *Rhizobium* strains were grown on solid and liquid LB media supplemented with 2.5 mM of MgSO_4_ and CaCl_2_ (LBmc) at 30°C. ST18 *E. coli* strains were grown on solid and liquid LB media supplemented with 5-aminolevulinic acid hydrochloride (ALA) 50 µg mL-1. The following concentrations of antibiotics were used when working with both *E. coli* strains ST18 and DH517 (μg·mL-1): chloramphenicol (Cm), kanamycin (Km) 25, and tetracycline (Tc) 5. When working with *R. leguminosarum, S. meliloti* or *S. medicae* strains, the following concentrations of antibiotics were used (μg·mL-1): neomycin (Nm) 200, streptomycin (Sm) 200, and tetracycline (Tc) 5.

**Table 1.**
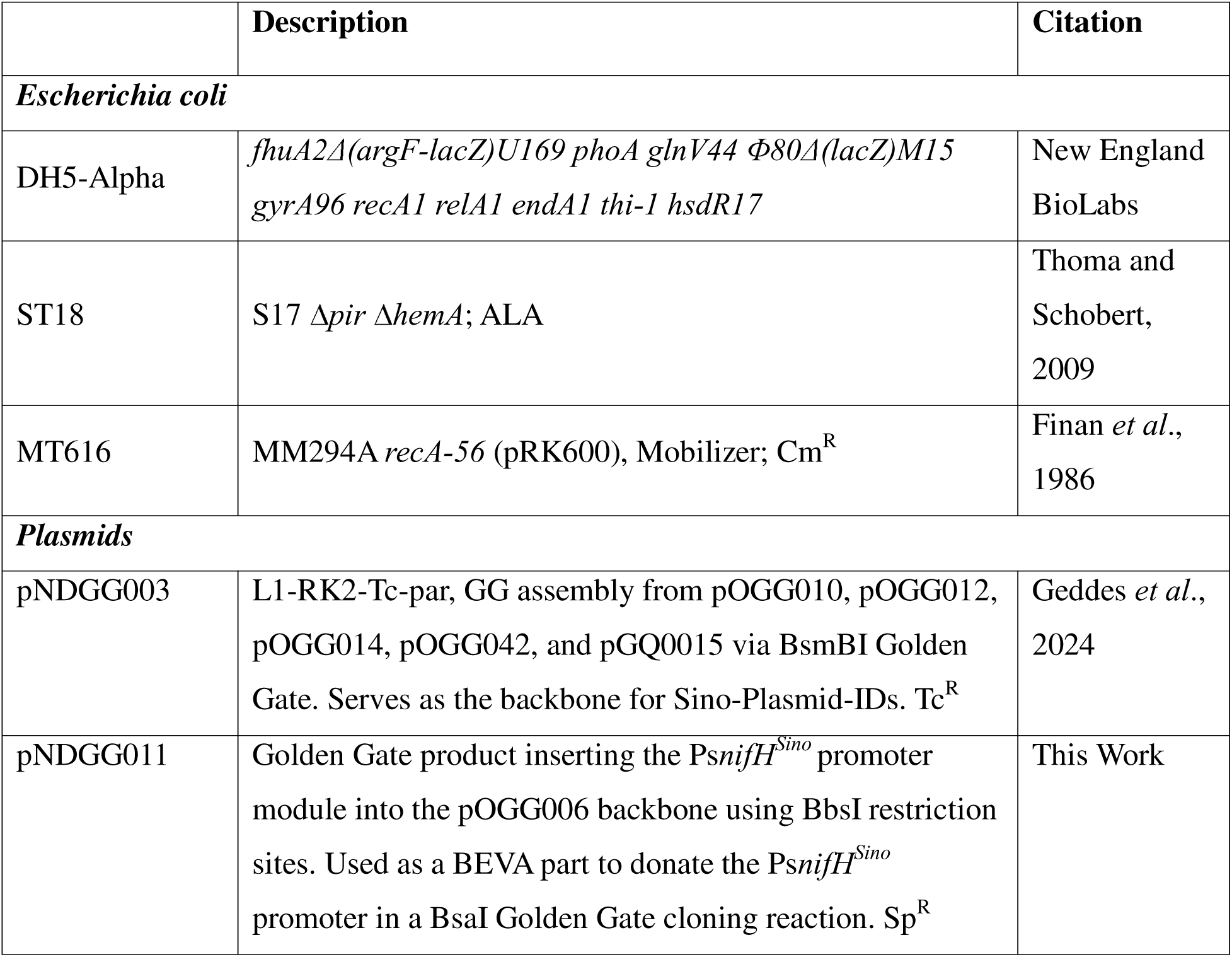
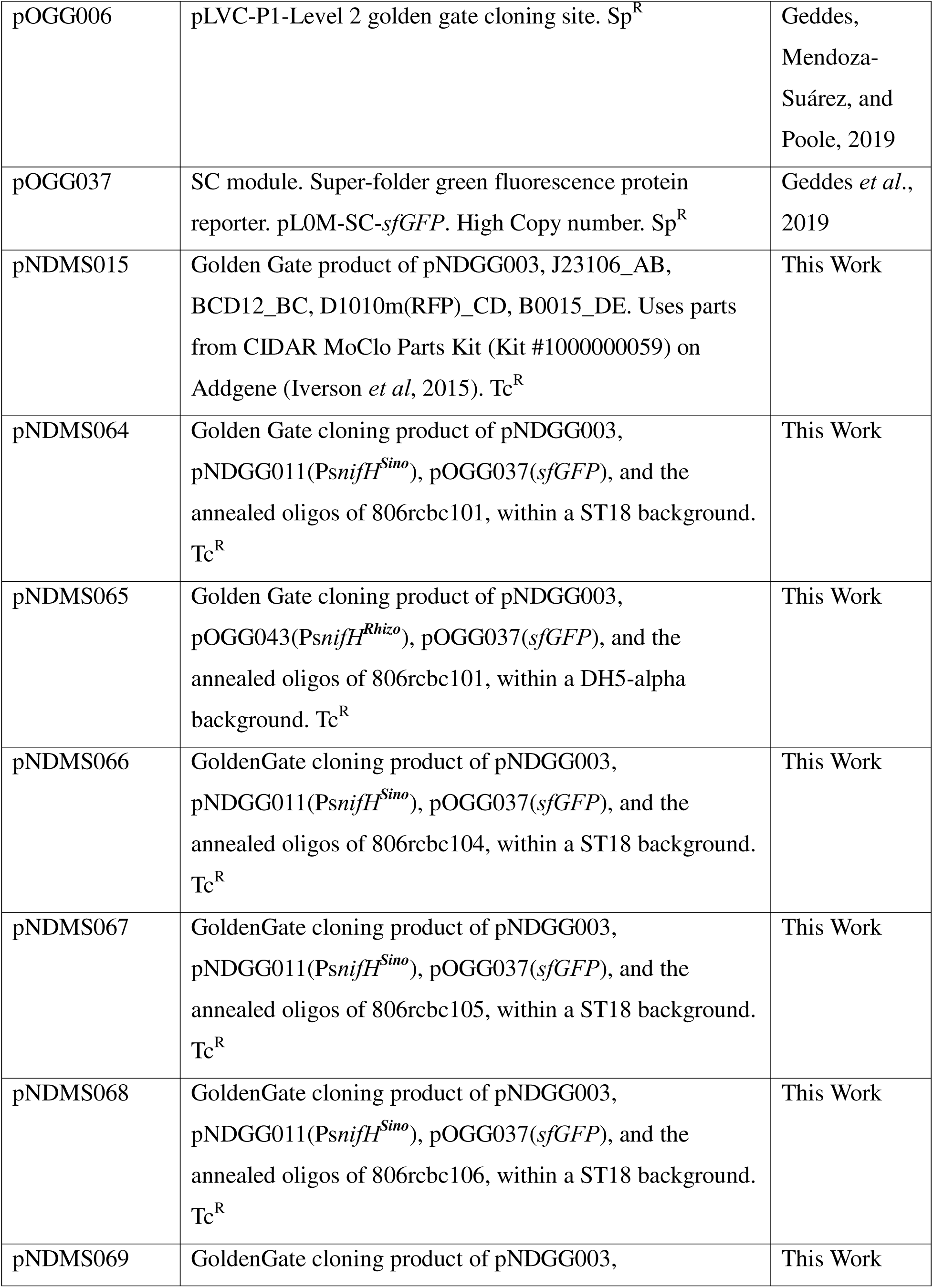
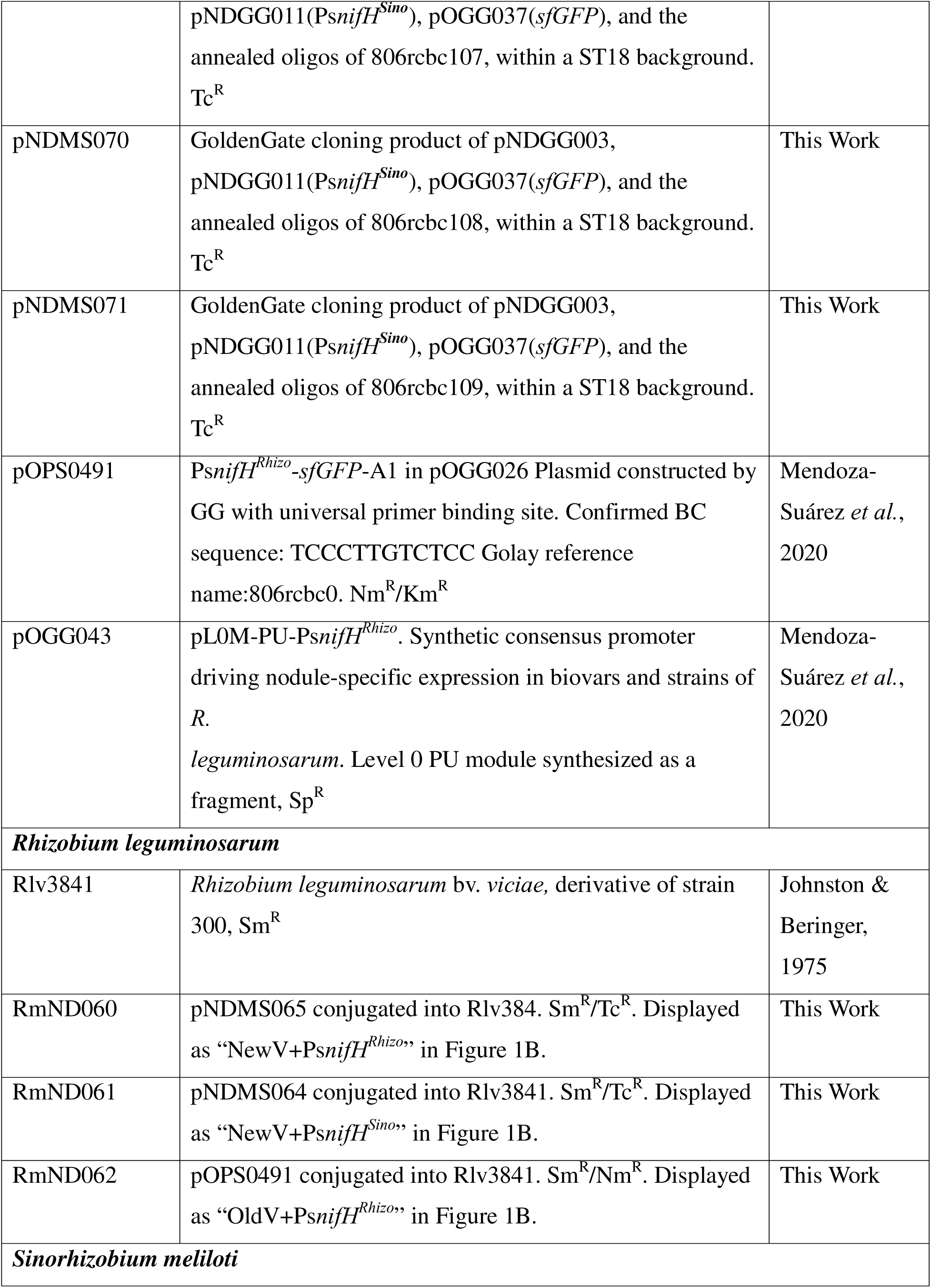
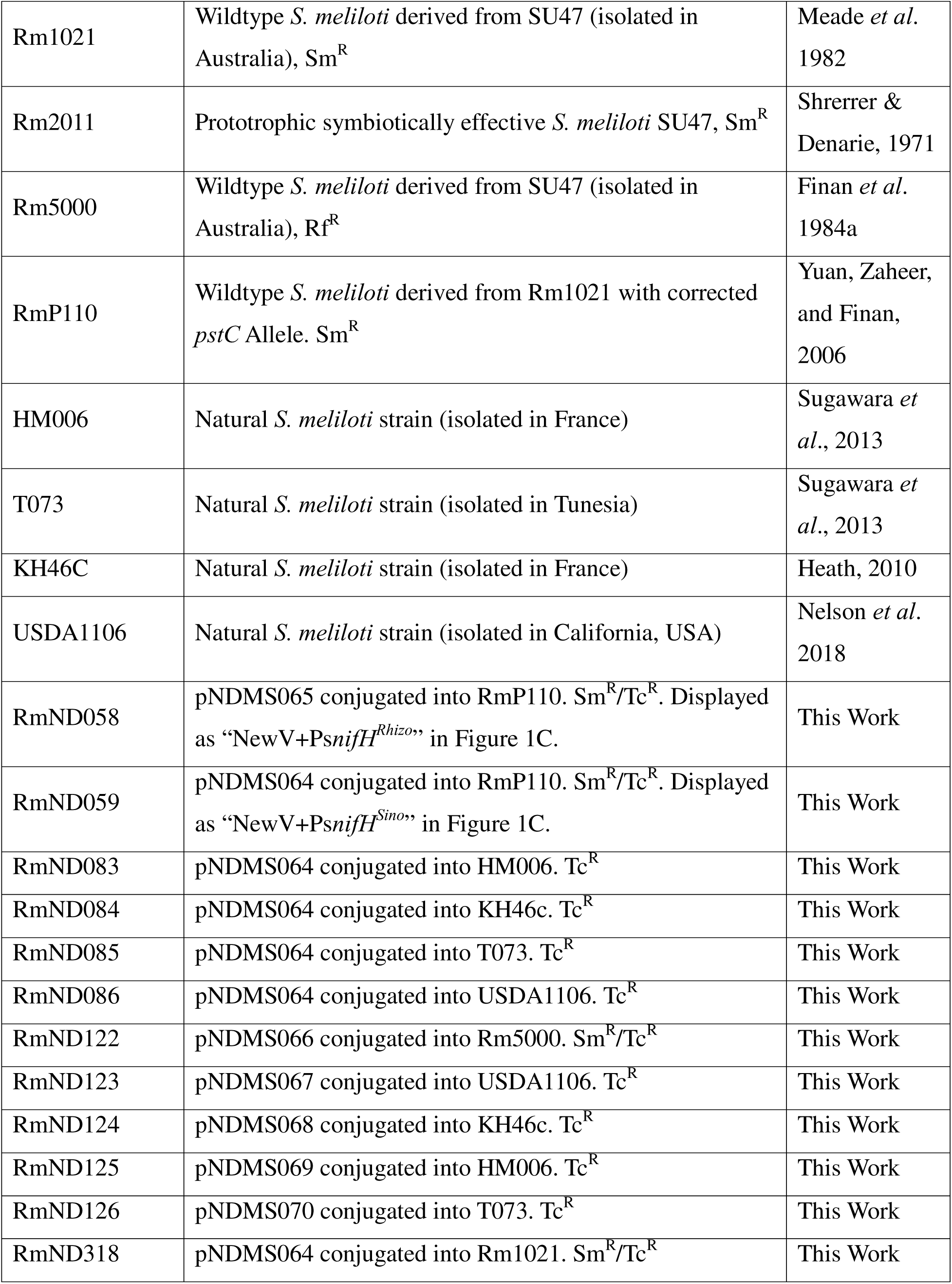
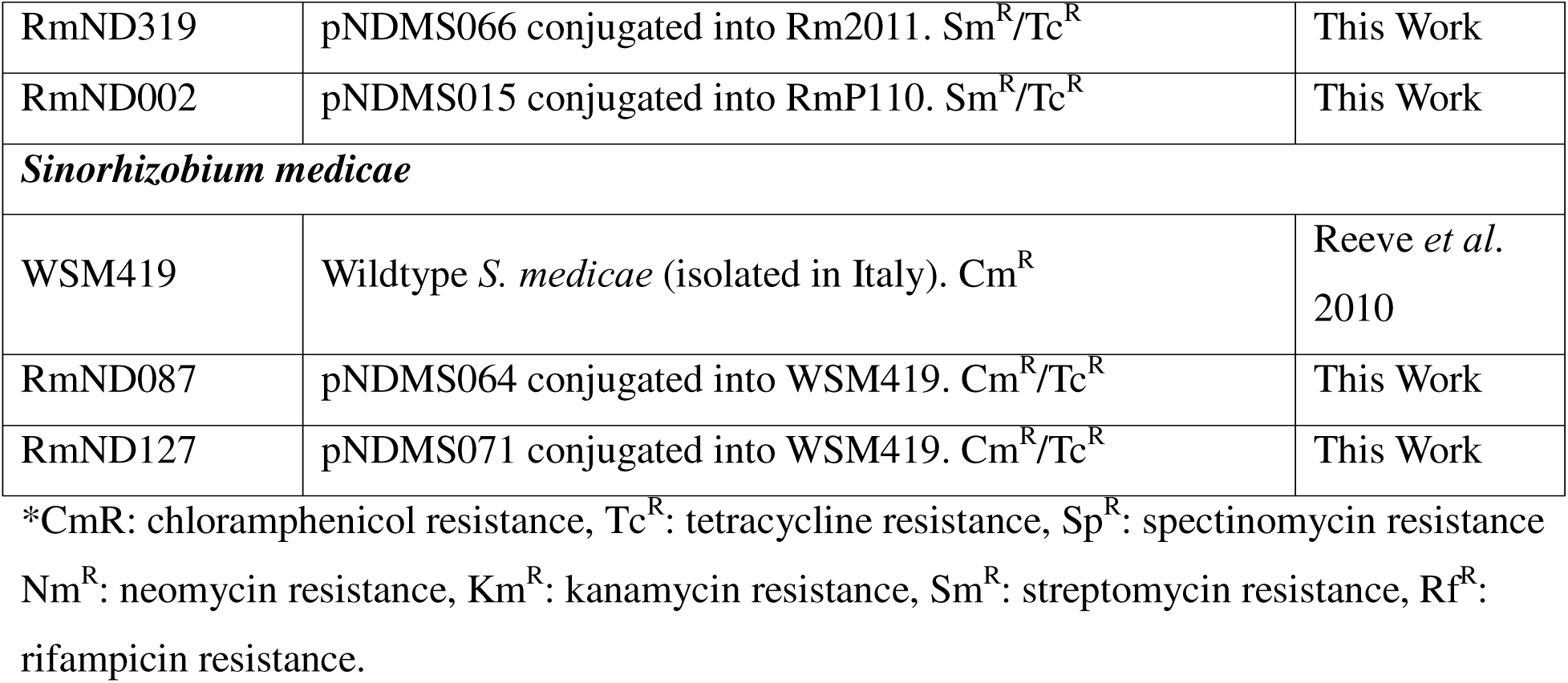
Bacterial Strains and Plasmids.

### Design of synthetic Ps*nifH^Sino^* consensus promoter

A 3110bp sequence of *nifH* and its upstream Intergenic Region (IGR) from *S. meliloti* 1021, (Sm1021*nifH*+IGR), was used as a template to find *Sinorhizobium* genomes in the Integrated Microbial Genomes & Microbiomes (IMG/M) system (Markowitz *et al*., 2012). As a result, 110 genomes (including draft genomes) were collected August 25, 2021. BLASTn (DNA vs. DNA E-value of 1e-5) was run on the 110 genomes. Scaffolds were generated of each strain and those with an E-value of 0 were extracted. Scaffolds with incomplete *nifHDK* operons were removed leaving 81 scaffolds with complete *nifHDK* operons. The resulting scaffolds included exclusively sequences from *S. meliloti* and *S. medicae* species. The 81 scaffolds were aligned in the bioinformatics software Geneious (Geneious Alignment using default settings). A 368 nt consensus sequence directly upstream of *nifH* was selected for the Ps*nifH^Sino^* promoter that includes a predicted NifA upstream activator sequence (UAS, nt 168 through 183), RpoN-binding site (nt 272 through 287), and ribosome binding site (RBS, nt 358 through 363), with a start codon ATG at nt 369 (Supplemental Figure SF1A). BsaI sites were added to both the 5’ and 3’ ends of the 368nt consensus sequence with GGAG and AATG cut-sites to act as a PU module in a Level 1 Golden Gate cloning (Weber *et al*., 2011). Additionally, BbsI sites were added to both the 5’ and 3’ ends of the PU module with TGCC and GGGA overhangs in order to clone the part into a pOGG006 backbone to generate a Ps*nifH^sino^* PU donating plasmid (pNDGG011) in DH517 *E coli.* The structure of the synthesized 420nt oligonucleotide before cloning into pOGG006: 5’-BbsI-BsaI-Ps*nifH^sino^*-BsaI-BbsI-3’ (Supplemental Figure SF1B).

### Sino-Plasmid-ID Construction

Level 1 Golden Gate cloning was used to create a library of unique Sino-Plasmid-IDs. Each reaction included the following components: T4 DNA ligase, BsaI restriction enzyme, backbone - pNDGG003, PU module - Ps*nifH^sino^* from pNDGG011, SC module – *sfGFP* from pOGG037, and annealed sense and antisense oligonucleotides containing a single unique Barcode-ID (806rcbc101, 806rcbc104-806rcbc109 (Mendoza-Suárez *et al*., 2024)) in a one-tube one-step reaction (Engler, Kandzia and Marillonnet, 2008). Seven Sino-Plasmid-IDs (pNDMS064, pNDMS066-pNDMS071) were used in the experiments in this study and listed in Table 1, but 87 more have been created and deposited in Addgene.

### Conjugating unique Sino-Plasmid-ID vectors into rhizobia

Sino-Plasmid-IDs housed within ST18 *E. coli* were routinely transferred into *S. meliloti* by conjugation via biparental mating. In brief, overnight cultures of each strain were combined and spotted onto LBmc+ALA then incubated overnight at 30°C. After 24 hours, mating spots were resuspended into 1mL of physiological saline (0.9% NaCl), then plated on selective LBmc+Tc at 10^-1^, 10^-2^, and 10^-3^ dilutions and incubated for three days at 30°C. Single colonies from dilution plates were purified by three rounds of streak-plating.

### Plant assay setup

Plant assays were conducted in Leonard jar assemblies with a 1:1 (wt:wt) mixture of sand and vermiculite. This sand-vermiculite mix was supplemented with 250ml per pot of either Jensen’s nitrogen-deficient medium (Vincent, 1970) for alfalfa plants, or Nitrogen-free rooting solution (Poole *et al*., 1994) for pea plants, and then autoclaved. Six germinating seeds were added to each Leonard jar and inoculated two days later with 10 mL inoculum containing the appropriate rhizobia strain/s. Seedlings and soil were kept sterile until the seedlings were inoculated.

Plants used in this experiment included *Medicago sativa* cv. Iroquois (alfalfa) and *Pisum sativum* cv. Striker (green field pea). Alfalfa seeds were surface sterilized for five minutes with 95% ethanol and for 20 minutes with 2.5% NaClO. Alfalfa seeds were then rinsed with sterile water for one hour, with the water replaced every 15 minutes. Pea seeds were surface sterilized for 30 seconds in 95% ethanol followed by 5 minutes in 2% NaClO, then rinsed 10x with sterile water. Alfalfa and pea seeds were placed on 0.8% water agar, placed in a dark drawer at room temperature, and allowed two days to germinate. Each replicate pot was inoculated with 10^6^ CFU total of rhizobia mixed with sterile water two days after planting the germinated alfalfa seeds. Competition experiments involving multiple strains within one inoculum were inoculated with equal ratios of each strain. Actual ratios in mixed inoculants were verified by NGS sequencing (Supplemental Table ST3). Water control pots were given 10 mL of sterile water instead of an inoculum. Mixed inoculum 1 = RmND059, RmND122-RmND127. Mixed inoculum 2 = RmND123-RmND127, RmND318, RmND319. Plants were grown in Conviron Gen1000 and Conviron Gen2000 growth chambers with LED lights. The chambers were programmed to have a day cycle of 18 hours at 21°C with maximum light followed by a night cycle of 6 hours at 17°C with no light. Pea plants were harvested 25-33 days post inoculation. Alfalfa plants were harvested 31-34 days post inoculation

### Data collection for rhizobial nitrogen-fixing effectiveness

#### sfGFP Fluorescence

Individual nodules were picked off plants into wells of a Greiner 96 Black Flat Bottom Fluotrac. sfGFP Analysis was performed using an Aglient Biotek Cytation 5 plate reader. sfGFP Monochrometer (F) Scan settings = Excitation: 485/20, Emission: 528/20, Optics position: Top, Gain: extended, Read Speed: Normal, Delay: 100msec, Measurements per data point: 10, Light Source: Xenon Flash, Lamp Energy: High, Dynamic Range: Extended, Read Height: 7.00 mm, Number of points: 7 x 7 equally spaced within the well, Point Spacing (microns): 825 825, well diameter: 6960, probe diameter: 2000.

#### Acetlyene reduction assay

Using an Agilent HP6890 gas chromatograph (GC) system, whole plant roots were measured via acetylene reduction rate using 14mL acetylene injections in 140mL bottles (Hardy *et al*., 1968; Zhang *et al*., 2012). Three acetylene/ethylene readings were taken per replicate, each 6 minutes apart to ensure linearity of acetylene reduction activity.

#### Shoot dry weight

After the growth period, plant shoots were cut off at the soil line and placed in a drying oven for 11-13 days, then weighed.

#### Nodule size analysis

Individual nodules within wells of a 96-well plate were imaged using an Agilent BioTek Cytation 5 Plate Reader. Adobe Photoshop software was used to extract an image of only the nodules (magic wand tool). The image of just the nodules was analyzed with ImageJ to determine the number of pixels (Schneider *et al*., 2012). ImageJ settings = image type: 8-bit, minimum threshold: 0, maximum threshold: 200. Nodule size in pixels was converted to nodule size in mm^2^ via comparison with the known width of wells in the plates used for imaging.

### Data collection for rhizobial competitiveness for root nodulation

#### DNA extraction from nodules

Depending on methodology, nodules were crushed individually in wells of a 96-well plate or crushed in a 1.5mL centrifuge tube as a pooled group with all nodules from a replicate. Alkaline PEG 200 was added (at 10x sample volume) to crushed nodules (Chomczynski and Rymaszewski, 2006). Samples were incubated for 15 min at 60°C and vortexed. Plant tissue was precipitated at 1000 rcf for 5 minutes and the supernatant was transferred to a new tube for use as template in library preparation PCR template.

#### Library preparation for multiplex Illumina MiSeq sequencing strategy

Two-step amplicon sequencing of Plasmid-IDs was performed according to the “16S Metagenomic Sequencing Library Preparation” protocol from Illumina. Primers that amplify Plasmid-IDs were used for the primary PCR, and included 4-6 nt barcode sequences in between 5’ Plasmid-ID binding sites and 3’ Illumina sequencing primer binding sites that anchored the secondary PCR (Supplemental Table ST1) (Mendoza-Suárez *et al*., 2024). A total of 12 unique forward Plasmid-ID primers were combined with 8 unique reverse primers to index the wells of a 96-well plate in the primary PCR. The secondary PCR used standard two-step PCR primers that added Nextera barcodes and Illumina adaptors (Supplemental Table ST2) (Gohl *et al*., 2016). Libraries were sequenced on Illumina MiSeq.

### Data manipulations, statistical analysis and figure preparation

Paired-end reads from Illumina MiSeq sequencing were merged using VSEARCH (Rognes *et al*., 2016). Merged reads were then deconvoluted into individual samples based on forward and reverse PCR primers, corresponding to rows and columns (respectively) of a 96-well plate.

Within each sample, plasmids were identified by Golay barcodes (Caporaso *et al*., 2012) mapped to a reference barcode database (stored as a .csv file). Both the primer deconvolution and barcode mapping were performed using a custom Python script, plasmid_ID.py, based on previous work (Mendoza-Suárez *et al*., 2020). The output from this step is a .csv file with the count of each identified barcode for each sample. Code and documentation for this study is available on GitHub (https://github.com/NDSU-Geddes-Lab/plasmid-id). The code contained in the GitHub repository is only for pre-processing the data; statistical analysis was performed *ad hoc*. Further data manipulation was done in Microsoft Excel to match the barcode IDs with the appropriate strain and background.

Statistical Analysis was performed using Prism GraphPad. All statistical analysis can be found in Supplemental Statistics SS1.

Figures and graphs were prepared using Prism GraphPad and BioRender. Plasmid maps and sequence alignments were prepared using Geneious.

## Results

### Re-design of Plasmid-IDs to function in *Sinorhizobia*

We focused on two aspects for re-design of Plasmid-IDs to function in *Sinorhizobia*. First, the Nm^R^-plasmid backbone pOGG026 from the Bacterial Expression Vector Archive (BEVA) was replaced with an analogous Tc^R^-based backbone, pNDGG003, from BEVA2.0 which showed more effective antibiotic selection in *S. meliloti* (Geddes, Mendoza-Suárez and Poole, 2019; Geddes *et al*., 2024). To further adapt the system for *Sinorhizobia*, we anticipated a need to refine the promoter part of the Plasmid-ID that drives sfGFP expression, which was designed based on a consensus of *Rhizobium* genus *nifH* promoter sequences (Mendoza-Suárez *et al*., 2020). In a similar workflow, we generated a synthetic consensus promoter for *Sinorhizobia* by aligning 81 sequences upstream of the *nifH* CDS from publicly available *S. meliloti* and *S. medicae* genomes (Supplemental Figure SF1A). The consensus sequence, 368 bp upstream of *nifH* was used as the new *Sinorhizobia* synthetic promoter we called Ps*nifH^Sino^* and included an upstream activator sequence for NifA (TGT-N_10_-ACA), a recognizable σ54 binding site (TGGCAC-N_5_-TTGCA), and a ribosome binding site (RBS) (AGGAGG). The Ps*nifH^Sino^* promoter was combined with an *sfGFP* cassette and oligonucleotides containing Golay barcodes into pNDGG003, conserving the previously reported Plasmid-ID architecture to generate new Sino-Plasmid-IDs (Mendoza-Suárez *et al*., 2020) (Figure 1A).

**Figure 1.**
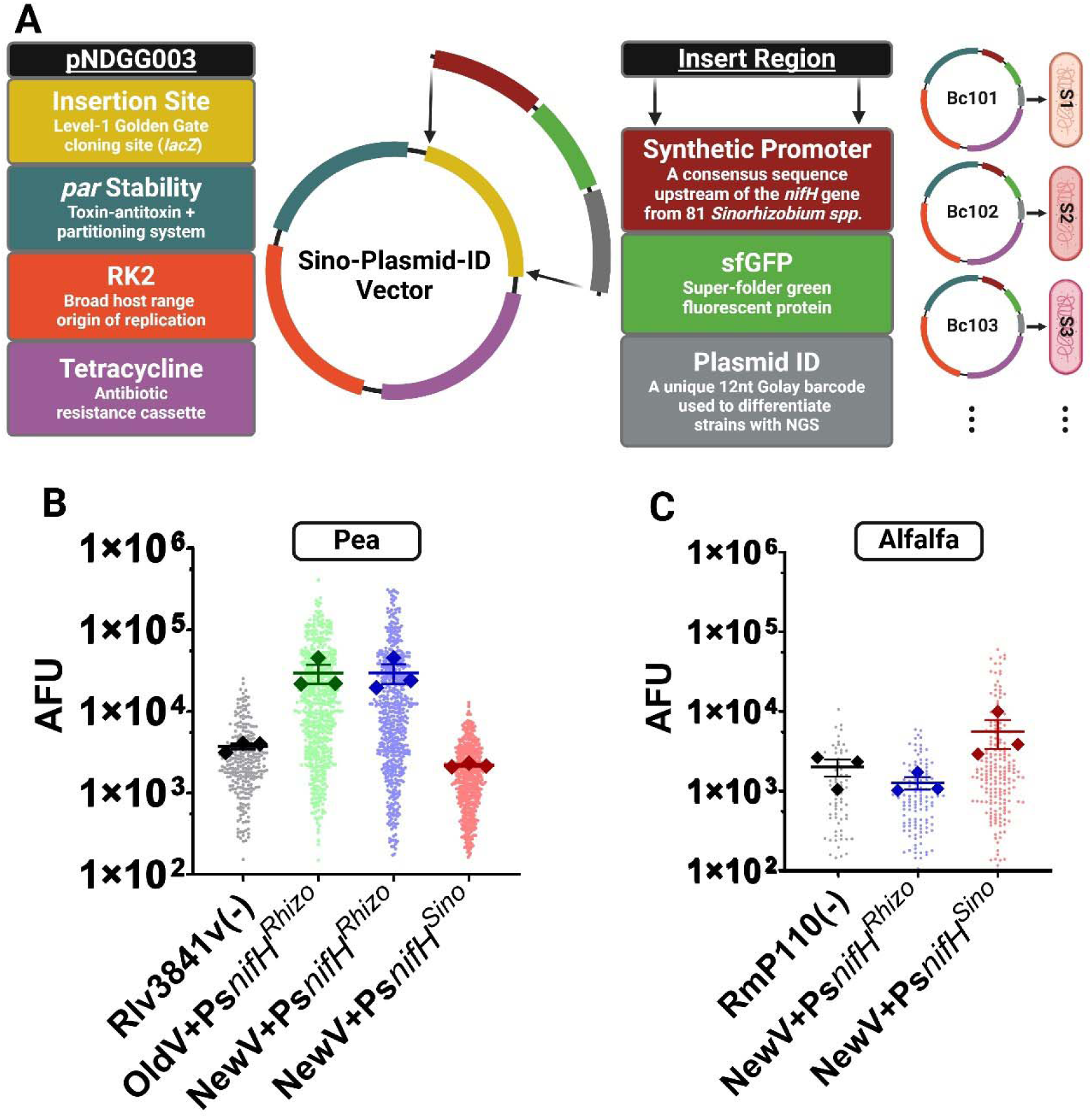
Sino-Plasmid-ID Design and comparison to Rhizo-Plasmid-ID. (A) Design of the *S. meliloti* Sino-Plasmid ID vector. The vector backbone (pNDGG003) includes a tetracycline resistance cassette, an RK2 origin of replication, partitioning genes and a toxin-antitoxin system (*parABCDE*), and a level 1 Golden Gate cloning site. Unique Sino-Plasmid-IDs were created by cloning an insert region into the level 1 Golden Gate Cloning site. The insert region included Psn*ifH^Sino^*, an *sfGFP* cassette, and a unique Plasmid ID containing a 12nt Golay barcode. These unique Sino-Plasmid-IDs were conjugated into separate strains of *Sinorhizobium* to enable tracking via NGS. (B) Displays nitrogen-fixing effectiveness via AFU intensity in *R. leguminsarum* strains (Rlv3841 background) in pea nodules while (C) displays strains of *S. meliloti* strains (RmP110 background) in alfalfa nodules. AFU measures the amount of sfGFP expression from individual nodules. Small circles represent all the individual nodules in all replicates. Large diamonds represent the mean AFU value from nodules in a replicate pot. Error bars represent SEM. (B) Rlv3841(-) = Rlv3841 = wildtype *R. leguminosarum* with no vector. OldV+Ps*nifH^Rhizo^* = RmND062 = former vector backbone (pOGG026) with Ps*nifH^Rhizo^*. NewV+Ps*nifH^Rhizo^* = RmND060 = updated vector backbone (pNDGG003) with Ps*nifH^Rhizo^*. NewV+Ps*nifH^Sino^* = RmND061 = updated vector backbone with Ps*nifH^Sino^*. (C) RmP110(-) = RmP110 is the wildtype strain of *S. meliloti* with no vector. NewV+Ps*nifH^Rhizo^* = RmND058 = updated vector backbone with Ps*nifH^Rhizo^* promoter. NewV+Ps*nifH^Sino^* = RmND059 = updated vector backbone with Ps*nifH^Sino^*.

### New Sino-Plasmid-IDs show stability in the absence of antibiotic selection

The stability of Plasmid-IDs is crucial because they need to be maintained in the absence of antibiotic selection during nodulation of the host. The stability of the new plasmid-ID architecture was verified in liquid culture and *in planta*. For liquid validation, we replaced the Ps*nifH^sino^*promoter with a constitutive promoter driving RFP production in the pNDGG003 backbone and introduced the resulting plasmid to *S. meliloti* RmP110. The resulting strain (RmND002) was grown for 24 hours (∼10 generations) in three independent replicates and plated by serial dilution. The number of fluorescent colonies was used to measure plasmid loss. In two of the three replicates, the plasmid was present in 100% of colonies even after 10 generations. In the third replicate only 0.128% of the colonies were non-fluorescent, indicating a loss of their Plasmid-ID. In all, only 3/7107 colonies were found to be non-fluorescent. Next, we introduced Sino-Plasmid-IDs with the Ps*nifH^Sino^*:sfGFP module into *S. meliloti* RmP110 and tested the stability of plasmids during host plant nodulation. In the *in planta* experiment, every single nodule (n=107 from 12 plants) showed the presence of their Sino-Plasmid-ID (via tetracycline resistance) after 33-35 days of plant growth, reinforcing evidence of stability in the absence of antibiotic selection.

### *Sinorhizobium* and *Rhizobium* synthetic *nifH* promoters are not cross-compatible

To investigate the performance of the new Ps*nifH^Sino^*promoter module, we introduced Sino-Plasmid-IDs to both *S. meliloti* (RmP110) and *R. leguminosarum* (Rlv3841). We also cloned the *Rhizobium* Ps*nifH* promoter (Ps*nifH^Rhizo^*) from the previous system into pNDGG003 in the same Plasmid-ID architecture to allow comparisons of the two promoters in *Sinorhizobia* (Rhizo-Plasmid-ID) and introduced this plasmid to each strain. These were further compared to a previously published plasmid in Rlv3841(pOPS0491) ((Mendoza-Suárez *et al*., 2020). These conjugated strains along with wildtype controls without Plasmid-IDs were measured for evaluation of sfGFP expression of nodules following isogenic inoculation of alfalfa or pea plants. Pea nodules formed by *R. leguminosarum* bearing Rhizo-Plasmid-IDs (RmND060) fluoresced similarly to those bearing the original Plasmid-IDs (RmND062). In stark contrast pea nodules formed by *R. leguminosarum* with Sino-Plasmid-IDs (RmND061) showed an absence of fluorescence closer to nodules formed by plasmid-less wildtype cells (Figure 1B, Supplemental Statistics SS1). Likewise, in alfalfa nodules formed by *S. meliloti,* the Sino-Plasmid-IDs resulted in a significant fluorescence signal (RmND059), whereas the Rhizo-Plasmid-IDs (RmND058) showed background levels of fluorescence (Figure 1C, Supplemental Statistics SS1). These data indicate that the *nifH* promoters of *Sinorhizobia* and *Rhizobia* are not cross-compatible.

### The Ps*nifH^Sino^*:sfGFP module functions as a biosensor of nitrogen-fixation effectiveness in *Sinorhizobia*

Next, we aimed to verify the utility of Sino-Plasmid-IDs to evaluate the effectiveness (ability to support the growth of their host plant via supplying bioavailable nitrogen) of rhizobia based on nodule fluorescence. Though this was previously validated for Plasmid-IDs in pea (Mendoza-Suárez *et al*., 2020), the overall fluorescence signal observed from *S. meliloti* Sino-Plasmid-IDs in alfalfa nodules was significantly lower than *R. leguminosarum* Plasmid-IDs in pea nodules (Figure 1 BC, Supplemental Statistic SS1). Using a test set of six *Sinorhizobium* strains with varying levels of nitrogen-fixing effectiveness (*S. meliloti* RmP110, USDA1106, T073, HM006 and KH46c, and *S. medicae* WSM419), we assessed whether Sino-Plasmid-IDs could accurately measure nitrogen-fixing effectiveness via sfGFP fluorescence of alfalfa nodules. Sino-Plasmid-IDs were conjugated into the six strains, and each was introduced as an isogenic inoculant to alfalfa plants. Following 32-34 days of growth, data for nitrogen-fixation effectiveness were collected by acetylene reduction assay and shoot dry weight, and the average fluorescence of all nodules from each replicate was measured. We observed significant correlations between nodule fluorescence (AFU) and both shoot dry weight (R^2^ = 0.7087) and acetylene reduction (R^2^ = 0.6938) indicating the Ps*nifH^Sino^*:sfGFP module of Sino-Plasmid-IDs functions effectively as a bioreporter of nitrogen-fixation in nodules (Figure 2, Supplemental Statistics SS1). We also evaluated the same correlations using nodule size or composite fluorescence/nodule size metrics, but found overall fluorescence to have the highest correlation (Supplemental Figure SF2, Supplemental Statistics SS1). Overall these data indicate the Ps*nifH^Sino^*:sfGFP bioreporter of Sino-Plasmid-IDs functions effectively as a biosensor of nitrogen-fixation activity in diverse *Sinorhizobium* strains.

**Figure 2.**
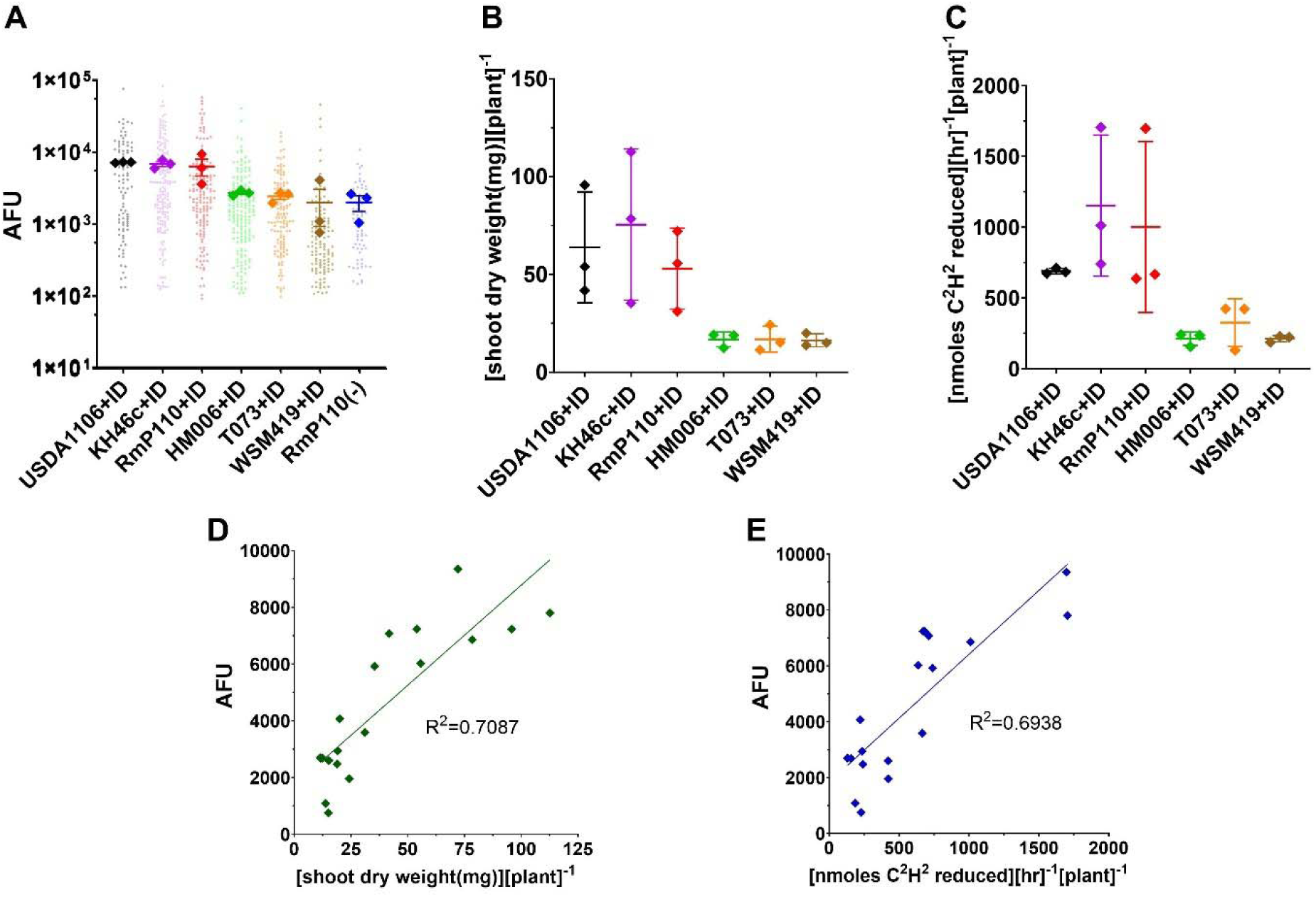
Validating the ability of Sino-Plasmid-ID to measure N_2_-fixing effectiveness. RmP110+ID, HM006+ID, KH46c+ID, T073+ID, USDA1106+ID, WSM419+ID = RmND059, RmND083, RmND084, RmND085, RmND086, RmND087 respectively and all contain the same Sino-Plasmid-ID. (A) Nitrogen-fixing effectiveness via intensity of AFU. AFU represents the amount of sfGFP expression. Small circles represent all individual nodules in all replicates. Large diamonds represent the mean AFU value from nodules in a single replicate pot. RmP110(-) represents the wild type RmP110 without a conjugated Sino-Plasmid-ID. Error bars represent SEM. (B) Nitrogen-fixing effectiveness via shoot dry weight. (C) Nitrogen-fixing effectiveness via acetylene reduction assay. (BC) Error bars represent standard deviation. (DE) Data points represent the replicate values of the 6 Sino-Plasmid-ID tagged strains in (ABC). (D) Correlation of sfGFP fluorescence with shoot dry weight. (E) Correlation of sfGFP fluorescence with acetylene reduction.

### Plasmid-ID barcodes allow evaluations of competitiveness for nodule occupancy in *Sinorhizobium* in individual or pooled nodules

Finally, we utilized the Sino-Plasmid-IDs to evaluate competitiveness for nodule occupancy and nitrogen-fixation effectiveness concurrently in our test strain set. This experiment included seven strains (the aforementioned six plus rifampicin resistant SU47 derivative Rm5000 (Finan *et al*., 1984b) labeled with unique Sino-Plasmid-IDs, co-inoculated onto alfalfa plants at equal cell densities. Following a similar workflow to Plasmid-ID, we assessed fluorescence and strain contents of individual nodules at 32 days post inoculation. We observed substantial differences in competitiveness for nodule occupancy among the strains, with KH46c dominating the nodule contents, and SU47 derivative “wildtype” strains RmP110 and Rm5000 rarely observed (Figure 3A, Supplemental Statistics SS1). We found nodule fluorescence intensity of certain strains differed between a mixed strain inoculum (Figure 3B) and a single strain inocula (Figure 2A).

**Figure 3.**
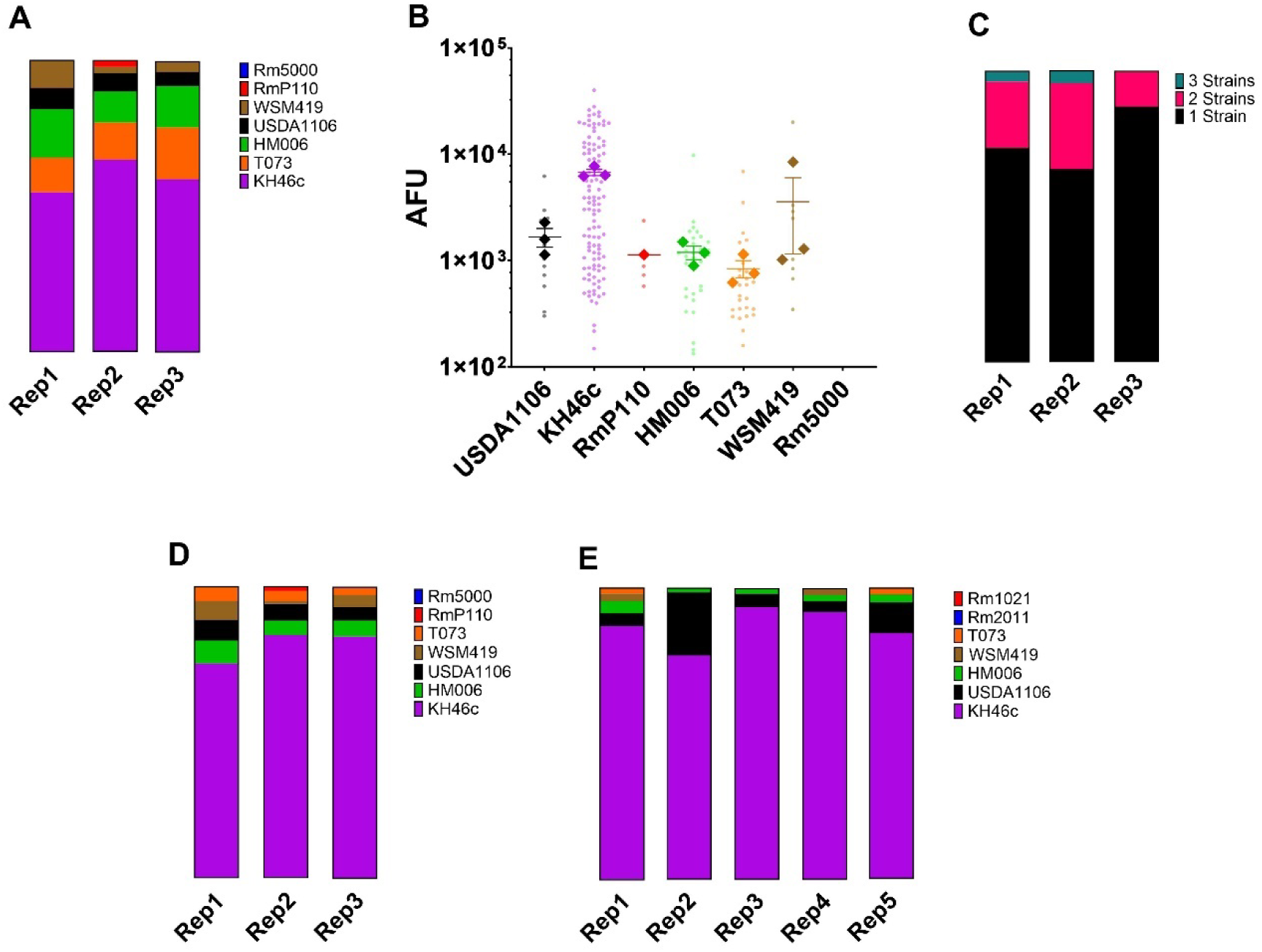
Validating the ability of Sino-Plasmid-ID to measure competitiveness for root nodulation. RmP110, Rm5000, USDA1106, KH46c, HM006, T073, WSM419, Rm1021, Rm2011 = RmND059, RmND122, RmND123, RmND124, RmND125, RmND126, RmND127, RmND318, and RmND319 respectively (A) Percentage of strains from total nodules occupied by individual nodule method. (B) Nitrogen-fixing effectiveness via AFU intensity in mixed inoculum. AFU represents the amount of sfGFP expression. Small circles represent all the individual nodules in all replicates. Large diamonds represent the mean AFU value from nodules in a replicate pot. Error bars represent SEM. (C) Frequency of mixed nodules. (D) Percentage of strains from total sequencing reads by pooled nodule method (mixed inoculum 1). (E) Percentage of strains from total sequencing reads by pooled nodule method (mixed inoculum 2).

KH46c nodules fluoresced relatively more intensely in mixed nodule scenarios, while USDA1106 and RmP110 saw a reduced intensity of sfGFP fluorescence (Figure 3B and Figure 2A). Similar to previously reported data in pea (Mendoza-Suárez *et al*., 2020), we were able to observe a high proportion of mixed nodules (24.1%), with some nodules even bearing 3 strains (2.6% of total) (Figure 3C).

Concurrently with the above experiment, we also evaluated the alternate utility of Plasmid-IDs to assess competitiveness using a pooled set of nodules from each replicate, saving significant time and cost (Figure 3D). In this scenario, competitiveness is assessed by the proportion of reads ascribed to a specific Sino-Plasmid-ID in the total nodule pool of the plant, rather than quantification of percent nodule occupancy. We generally found alignment of the nodulation competitiveness data using either approach, with KH46c dominating the population and Rmp110/Rm5000 nearly absent (Figure 3AD).

To more broadly explore the lack of competitiveness from lab domesticated SU47 derivative wild-type strains, we repeated the experiment replacing RmP110 and Rm5000 with Rm1021 and Rm2011, and similarly found the two wild-type strains to be nearly undetectable (Figure 3E). Overall, our test experiments support the utility of Sino-Plasmid-IDs for investigating competitiveness for nodule occupancy in *Sinorhizobium* strains.

## Discussion

In this work we describe the successful adaptation of the Plasmid-ID tool for measuring competitiveness for nodule occupancy and effectiveness of symbiotic nitrogen fixation to the *Sinorhizobium-Medicago* system. These two traits are not genetically linked in rhizobia, yet both traits are crucial to optimal symbiotic nitrogen fixation by a rhizobium inoculant strain deployed in agriculture (Mendoza-Suárez *et al*., 2021). Thus, while several techniques have emerged that utilize NGS to quantify rhizobium fitness and plant colonization, the Plasmid-ID system technique stands out for its unique flexibility of utility (Mendoza-Suárez *et al*., 2020). The system was recently utilized to provide key insights into how rhizobium strain diversity affects plant growth in agricultural scenarios by utilizing the high-throughput nature of the system to screen 399 *R. leguminosarum* strain interactions with 212 faba bean genotypes (Mendoza-Suárez *et al*., 2024).

Crucial to the successful adaptation from *Rhizobium* to *Sinorhizobium* was altering the *nifH* promoter used to drive sfGFP expression. *Sinorhizobium* and *Rhizobium nifH* consensus promoters were found not to be cross-compatible (Figure 1BC). This was surprising given the conserved mechanism for *nifH* gene expression in rhizobia and previous reports of successful heterologous *nifH* expression (Cebolla, Ruiz-Berraquero and Palomares, 1994; Dixon and Kahn, 2004). Though the overall fluorescence signal from alfalfa nodules was not as high as that in pea nodules, we observed very similar correlations between nodule fluorescence and nitrogen fixation in the *Sinorhizobium-*alfalfa system to those observed in the *R. leguminosarum* bv. *viciae* and peas (Mendoza-Suárez *et al*., 2020) (Figure 2DE). Interestingly, we observed a discordance between nodule fluorescence from *Sinorhizobium* strains in isogenic inoculation compared to mixed inoculation scenarios (Figure 2A and Figure 3B). The differences in fluorescence likely reflect plant sanctioning responses during mixed inoculation, highlighting the utility of symbiosis biosensors such as these for investigating resource allocation during symbiosis (Westhoek *et al*., 2021). However, it should be cautioned that comparative measurements of nodule fluorescence across strain collections were challenging in mixed scenarios, due to the exclusion of uncompetitive strains from nodules limiting dataset size (Figure 3B).

Using the Sino-Plasmid-IDs, we successfully differentiated highly competitive *Sinorhizobium* strains from non-competitive strains in two different experimental workflows. Sequencing individual nodule contents as described in the original Plasmid-ID system provided accurate nodule occupancy quantification, facilitated concurrent measurements of nodule fluorescence, and allowed us to identify mixed nodules with up to three strains (Figure 3C). However, this approach is also cost and labor-intensive due to the need to prepare sequencing libraries from each nodule. Alternatively, sequencing Plasmid-IDs from pooled nodule preps minimized costs and labor, but lacks the advantages described above and involves more convoluted data interpretation where the proportion of reads of a given strain reflects both the number of nodules occupied as well as the population size in those nodules. Regardless of approach, KH46c was found to be highly competitive, consistent with data reported from other groups (Bellabarba *et al*., 2021; Burghardt *et al*., 2022). We also found four wildtype strains derived from SU47, wherein the bulk of genetic study on *Sinorhizobium* has been done (Rm1021, Rm2011, RmP110 and Rm5000), to be very uncompetitive for nodule occupancy (Figure 3ADE). This emphasizes a need to explore the rhizobium pangenome to identify genetic determinants of competitiveness for nodule occupancy and reflects a disadvantage in focusing heavily on lab-domesticated isolates for our knowledge of genetics, despite the optimized genetic platforms they present.

Overall, we anticipate the development of the Sino-Plasmid-ID system to be a valuable tool for the research community, given the status of the *Sinorhizobium-Medicago* system as a central model for studying rhizobium-legume symbioses. The system, including 87 additional sequence verified Sino-Plasmid-IDs, has been deposited into Addgene to facilitate access.

## Acknowledgements

This work was supported by a New Innovator in Food & Agricultural Research (FFAR) grant to B. A. Geddes ID: FF-NIA21-0000000061m and a National Alfalfa & Forage Alliance alfalfa checkoff award to B. A. Geddes. This work used resources of the Center for Computationally Assisted Science and Technology (CCAST) at North Dakota State University, which were made possible in part by NSF MRI Award No. 2019077. We thank Megan Ramsett for research and logistical support throughout the project. We also thank Geddes lab members (Department of Microbiological Sciences, North Dakota University) for their assistance in harvesting and processing plant experiments.

## Competing Interests

Barney A. Geddes is a co-founder of Frontier Bioforge LLC. and Lilac Agriculture Inc.

## Supplemental Information

**Supplemental Figure SF1.**
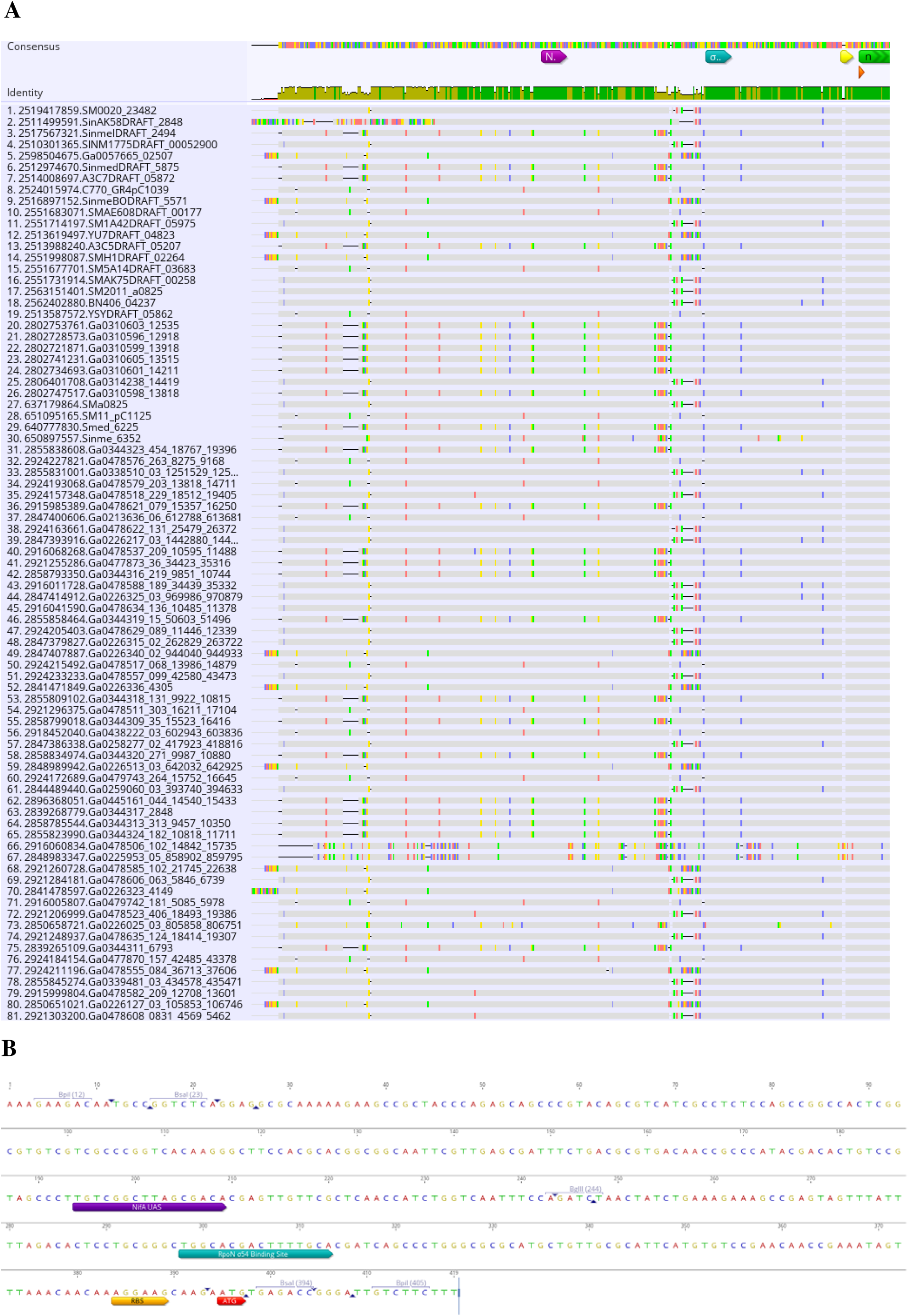
Consensus alignment of synthetic *nifH* promoter for alfalfa symbionts and module for cloning. (A) Alignment of 81 sequences upstream from the *nifH* gene forming a 368 nt consensus promoter. Includes a nifA upstream activator sequence (UAS) (TGT-N_10_-ACA) (purple), a binding sequence for RpoN σ^54^ (sigma factor for expression of all *nif* genes in nodules) (TGGCAC-N_5_-TTGCA) (blue), a ribosome-binding site (RBS) (AGGAGG) (yellow), and a start codon (ATG) (red). (B) Full Ps*nifH^Sino^* module design. Module was cloned into pOGG006 to create pNDGG011. Full PU module sequence: 5’- AAAGAAGACAATGCCGGTCTCAGGAGGCGCAAAAAGAAGCCGCTACCCAGAGCAG CCCGTACAGCGTCATCGCCTCTCCAGCCGGCCACTCGGCGTGTCGTCGCCCGGTCAC AAGGGCTTCCACGCACGGCGGCAATTCGTTGAGCGATTTCTGACGCGTGACAACCG CCCATACGACACTGTCCGTAGCCCTTGTCGGCTTAGCGACACGAGTTGTTCGCTCAA CCATCTGGTCAATTTCCAGATCTAACTATCTGAAAGAAAGCCGAGTAGTTTATTTTA GACACTCCTGCGGGCTGGCACGACTTTTGCACGATCAGCCCTGGGCGCGCATGCTGT TGCGCATTCATGTGTCCGAACAACCGAAATAGTTTAAACAACAAAGGAAGCAAGAA TGTGAGACCGGGATTGTCTTCTTT-3’

**Supplemental Figure SF2.**
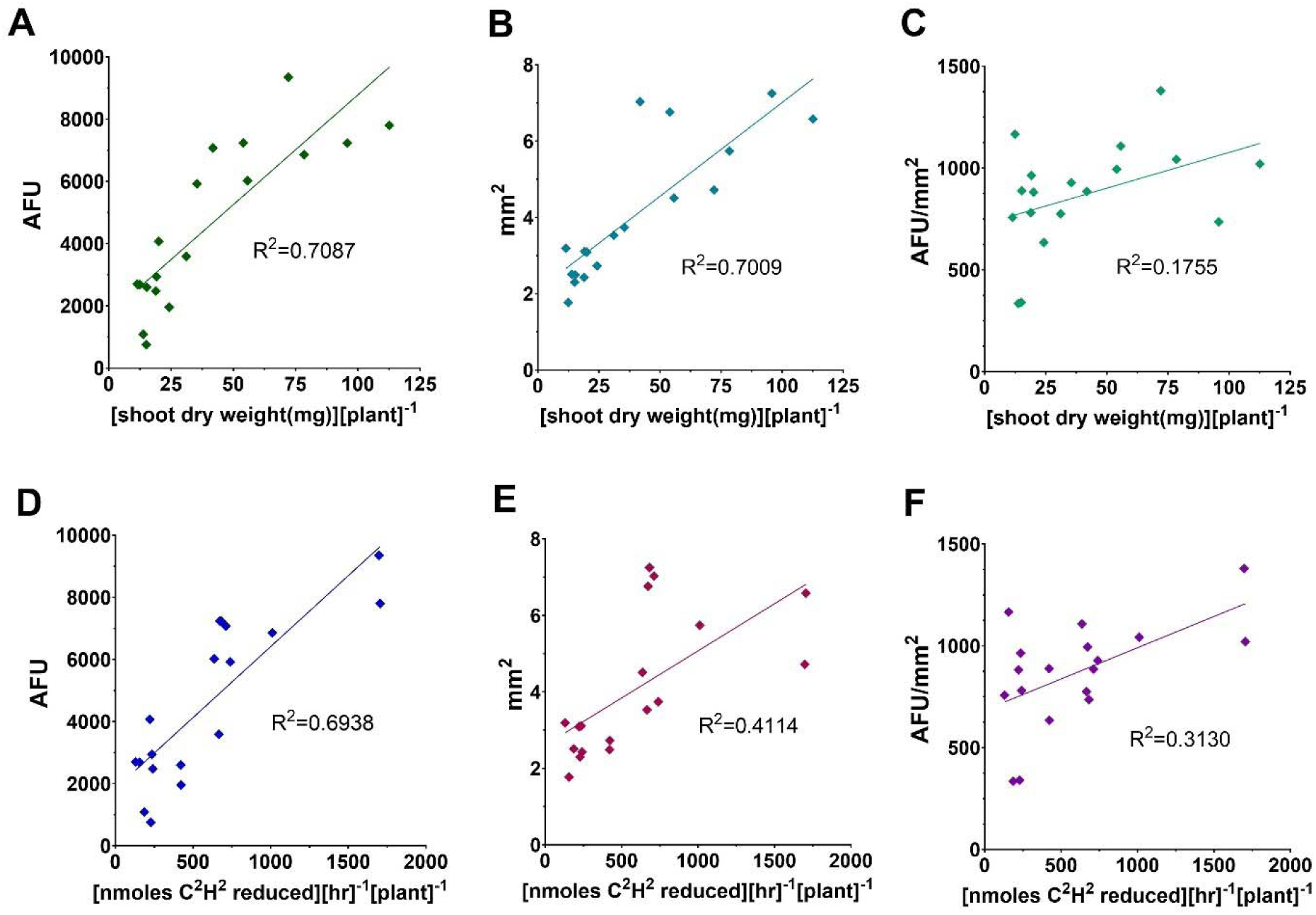
Comparing different metrics for measuring nitrogen-fixing effectiveness. This graph shows the correlation of three potential metrics for measuring nitrogen-fixing effectiveness (AFU(AD), mm^2^(BE), and AFU/mm^2^(CF)) against established measurements of nitrogen-fixing effectiveness: shoot dry weight (ABC) and acetylene reduction using gas chromatography (DEF). AFU represents the amount of sfGFP expression. Each data point represents a replicate value of RmND059, RmND083-RmND087 for the metrics displayed in the respective graph.

**Supplemental Table ST1.**
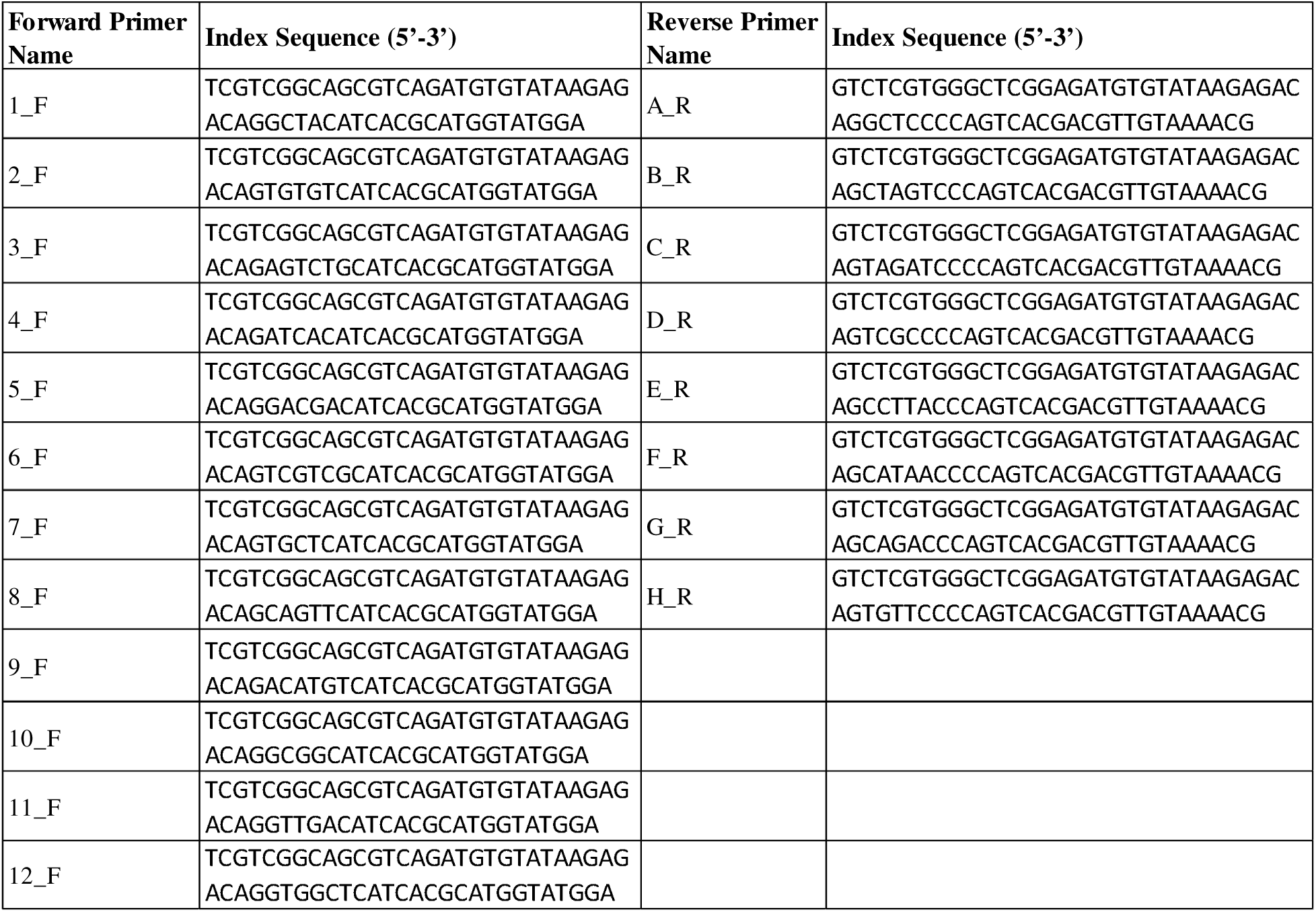
Forward and Reverse Primers for Primary PCR in library preparation. Primers displayed 5’ to 3’.

**Supplemental Table ST2.**
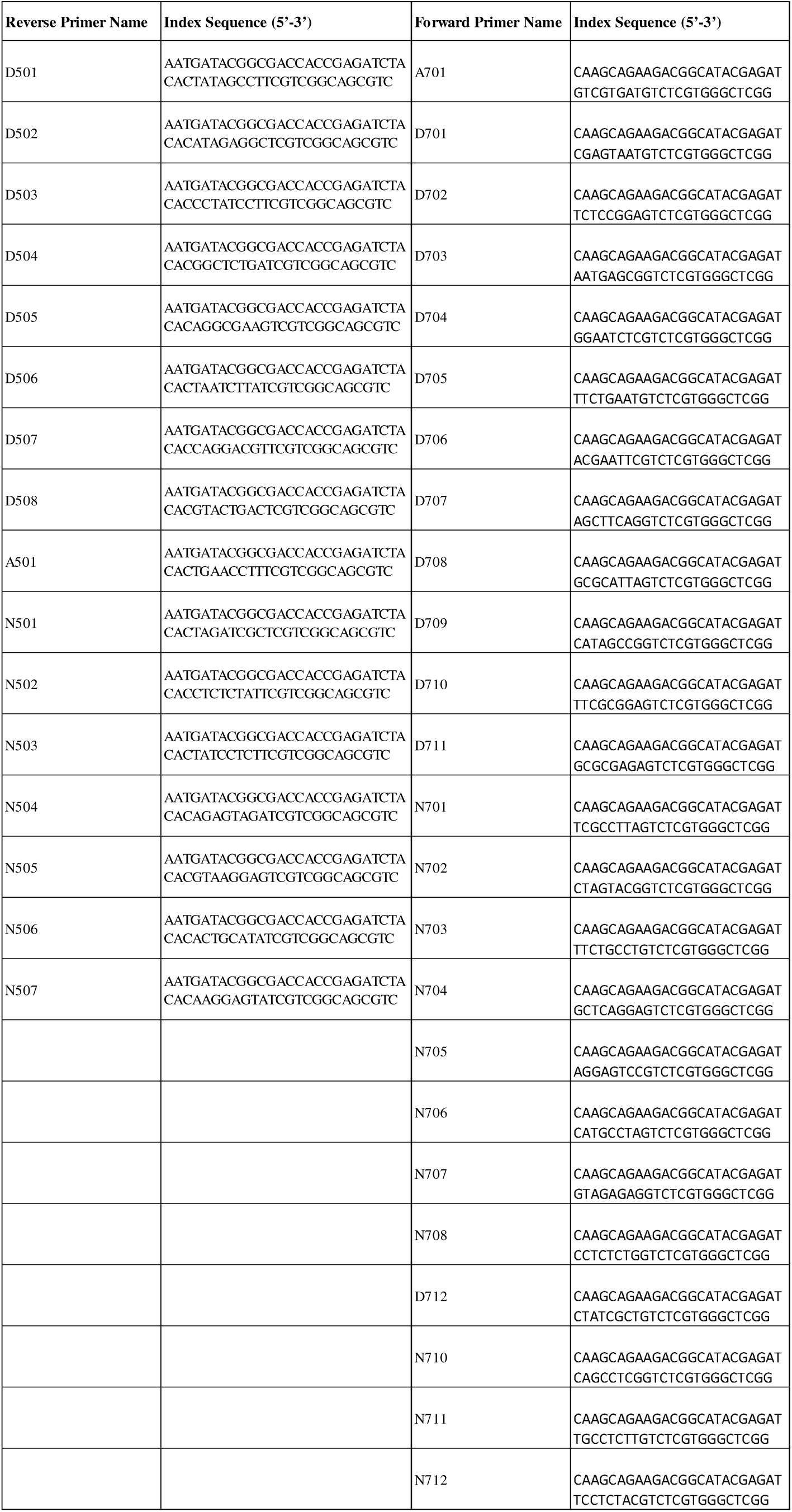
Forward and Reverse Primers for Secondary PCR in library preparation. Primers displayed 5’ to 3’.

**Supplemental Note SN1 – Modifications of Library Preparation Protocol from Illumina’s “16S Metagenomic Sequencing Library Preparation.”**

1. Primary PCR (“Amplicon PCR”):

a. The annealing temperature was increased to 64°C.
b. If processing using the pooled nodules method, then the template DNA was diluted 1/10 in ultrapure water and the cycle number was increased to 26 cycles.
2. Primary PCR Clean-Up (“PCR Clean-Up”): A 3:2 (Ampure XP beads : PCR) ratio was used.
3. Secondary PCR (“Index PCR”): The cycle number was increased to 12 cycles.
4. Secondary PCR Clean-Up (“PCR Clean-Up 2”): A 3:2 (Ampure XP beads : PCR) ratio was used.

**Supplemental Table ST3.**
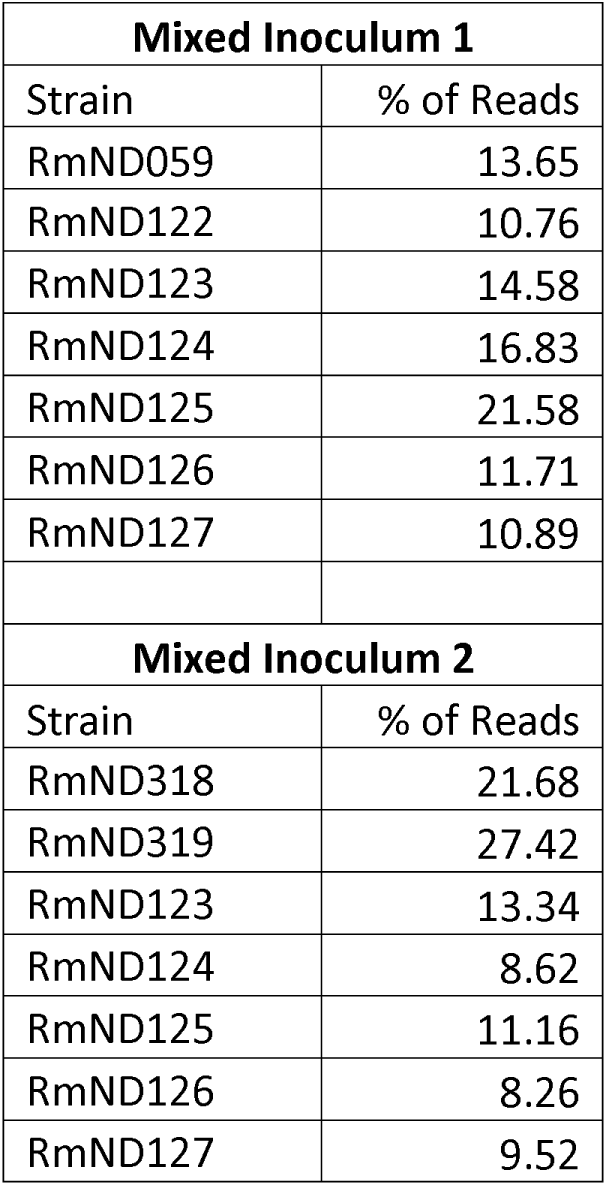
Composition of Plant Assay Inocula as verified by NGS of Plasmid-IDs. In competition experiments, each strain was intended to be inoculated with an equal number of CFU. The actual input ratios did not differ by more than 3.31x (RmND319 and RmND126 in mixed inoculum 2).

**Supplemental Statistics SS1 – Stats Excel Sheet**

Multiple sheets. Check all sheets.

## Notes

### Competing Interest Statement

Barney Geddes is cofounder of Lilac Agriculture Inc

